# Spontaneous axon regeneration is preserved despite gut microbiota disruption after spinal cord injury in lampreys

**DOI:** 10.64898/2026.02.06.704306

**Authors:** Laura González-Llera, Gabriel N. Santos-Durán, Ana Vences, Noemí Buján, Miguel Balado, Antón Barreiro-Iglesias

## Abstract

Traumatic spinal cord injuries (SCIs) often result in permanent disabilities in humans. One major reason for the lack of recovery is the inability of adult mammalian descending neurons to regenerate their axons after injury. In contrast, several fish species, such as the sea lamprey, exhibit spontaneous axon regeneration and successful functional recovery following a complete SCI. Recent studies have shown that a SCI in rodents and humans induces gut microbiome dysbiosis, which can impair recovery. Therefore, our goal was to examine how the microbiome changes after SCI in a regenerating animal model (the larval sea lamprey) and whether these changes influence the spontaneous regeneration of descending neuropeptidergic (cholecystokinergic) axons. Our data show that a complete SCI triggers an initial shift (5 weeks post-injury) in gut microbial communities in larval lampreys, characterized by an expansion of *Legionellaceae* family members. However, a treatment with broad-spectrum antibiotic gentamicin during the first 5 weeks post-injury, which completely disrupted the gut microbiome (eliminating *Legionellaceae* and promoting *Bradyrhizobiaceae* expansion), did not affect the spontaneous regeneration of descending cholecystokinergic axons at 10 weeks post-injury. This finding indicates that changes in the intestinal microbial communities following a complete SCI probably do not influence the spontaneous regeneration of descending axons in lampreys.

## 1 Introduction

Traumatic spinal cord injury (SCI) is a devastating event that in humans leads to permanent functional impairments, which includes permanent sensory, motor and autonomic impairments (Kim et al., 2023; Hosseini et al., 2024). The incidence of SCI is 10.5 cases per 100,000 people (over 700,000 new cases per year) (Kumar et al., 2018). At the cellular level, the lack of spontaneous recovery after a SCI can be related, among other causes, to the inability of axotomized neurons to regenerate their axons and reconnect with their appropriate targets above and below the injury site. Both intrinsic and extrinsic factors are known to limit axon regrowth and synaptogenesis after a SCI and to date, no effective treatment has been translated to the clinic to promote meaningful axon regeneration and neurological recovery after a SCI (see Zheng and Tuszynski, 2023). Thus, there is pressing need to deepen in our understanding of the cellular and molecular mechanisms limiting axon regeneration in the central nervous system (CNS) following a traumatic SCI.

As expected, there is a major focus into analysing the role that local CNS cells or of cells migrating to the site of injury (e.g., immune cells) play in regulating axon regeneration after SCI. However, SCIs are a whole-body disorder (Sauerbeck et al., 2015), and we should consider the role that peripheral systems/cells play in limiting or promoting recovery from SCI. The gut microbiome has been shown to play a role in nervous system development, in neurological disorders and in neuronal de/regeneration in different contexts (Kelly et al., 2016; Warner, 2019; Needham et al., 2020; Zeng et al., 2020; Iliodromiti et al., 2023; Loh et al., 2024). In rodents and humans, SCI causes gut microbiome dysbiosis (i.e., changes in gut microbial communities), which impairs recovery (Kigerl et al., 2016; Jogia and Ruitenberg, 2020; Du et al., 2021; Jing et al., 2021, 2023; Kong et al., 2023; Wilson et al., 2023). Moreover, recent work has shown that certain bacteria or bacteria derived metabolites can promote axon regeneration in the mammalian peripheral nervous system (Serger et al., 2022) or retina (Zhao et al., 2023).

In contrast to mammals, several fish species, like the sea lamprey, can recover from a complete transection SCI and regain most of their swimming capabilities (Katz et al., 2020; Fies et al., 2021; González-Llera et al., 2024; Tendolkar and Mokalled, 2025). In mature larval sea lampreys, recovery from SCI involves the regeneration of descending axons coming from the brainstem and synapse regeneration between regenerated axons and neurons below the site of injury (Jacobs et al., 1997; Cornide-Petronio et al., 2011; Parker, 2022; González-Llera et al., 2022, 2024). For example, we have recently shown that descending cholecystokinergic axons regenerate spontaneously after a complete SCI in larval lampreys, that these regenerated axons re-establish synaptic contacts below the site of injury and that swimming recovery correlates with the degree of regeneration of cholecystokinergic axons (González-Llera et al., 2024). The lamprey model of spontaneous regeneration after SCI has provided important knowledge on the role that different spinal cord intrinsic signalling systems play in promoting spontaneous regeneration (e.g., Lau et al., 2013; Fogerson et al., 2016; Romaus-Sanjurjo et al., 2018, 2019; Sobrido-Cameán et al., 2018, 2019, 2020a, 2020b; Rodemer et al., 2020; Sobrido-Cameán and Barreiro-Iglesias, 2022; Hu et al., 2017, 2021, 2023). However, there is not much knowledge on how the gut microbiome changes after SCI in larval sea lampreys or how those changes might affect spontaneous axon regeneration. For example, certain bacteria produce neurotransmitters (e.g., GABA: Han et al., 2023; Belelli et al., 2024; Tamés et al., 2024; serotonin: Sanidad et al., 2024). Interestingly, our work has shown that these neurotransmitters regulate axon regeneration after SCI in lampreys (GABA: Romaus-Sanjurjo et al., 2018, Sobrido-Cameán et al., 2018; serotonin: Sobrido-Cameán et al., 2019). Recent work has revealed the composition of the gut microbiome in larval sea lampreys (Tetlock et al., 2012; Mathai et al., 2021). Thus, we are now in an excellent position to analyse how the gut microbiome changes after a SCI in a regenerating animal like the larval sea lamprey and if these changes affect spontaneous axon regeneration in the spinal cord.

Here, we first studied the changes that occur in the composition of gut microbiome communities of larval sea lamprey after a complete transection SCI. Then, we studied how the gut microbiome is affected in lampreys with a SCI by a treatment with the broad-spectrum antibiotic gentamicin and if these changes affect the spontaneous regeneration of descending cholecystokinergic axons. Surprisingly, the gentamicin-induced changes in gut microbial communities did not affect the spontaneous regeneration of cholecystokinergic descending axons in injured lampreys.

## 2 Method

### 2.1 Animals and surgeries

Animal experimental procedures were performed in accordance with European Union and Spanish guidelines on animal care and experimentation and were approved by the Committee of Bioethics at the Universidade de Santiago de Compostela and the Xunta de Galicia government (Galicia, Spain; license number 15012/2020/011). Mature larval sea lampreys, *Petromyzon marinus* L. (n = 62; more than 90 mm in body length, 5 to 7 years of age), were used in this study. Larval lampreys were collected from the river Ulla (Galicia, Spain) with permission from the Xunta de Galicia and kept in an aerated freshwater aquarium (at 14ºC and fed with baker’s yeast) until they were used for the experimental procedures.

For the experiments, larvae were randomly assigned to the control non-injured, sham control or SCI (injured animals with or without gentamicin treatment; see below) groups. Animals were deeply anesthetized with 0.1% tricaine methanesulfonate (MS-222; Sigma) in lamprey Ringer solution (137 mM NaCl, 2.9 mM KCl, 2.1 mM CaCl2, 2 mM HEPES; pH 7.4) before surgical procedures and euthanized by decapitation (at the level of the 6th gill) at the end of the experiments (5, 8 or 10 weeks after surgery).

SCI surgeries were performed as described (Barreiro-Iglesias et al., 2014). Briefly, the spinal cord was exposed dorsally at the level of the gill area by making a longitudinal incision in the midline with a #11 scalpel. The complete spinal cord transection was performed at the level of the 5^th^ gill opening with Spring scissors (Fine Science Tools; Cat#15024-10). Completeness of the transection SCI was confirmed under the stereomicroscope by visualizing the spinal cord cut ends right after surgery and was confirmed 24 h later by checking that there were no caudally propagating movements below the site of injury. After surgery, larvae were put on ice for 1 h to allow the wound to dry and then allowed to recover in individual tanks with 200 ml of freshwater at 19ºC for 5, 8 or 10 weeks. Wounds are allowed to heal naturally, and no post-surgical wound care procedures are needed during recovery. During recovery/treatments, larvae were fed with 0.5 µL of baker’s yeast (60 mg/ml) 3 times per week (on Mondays, Wednesdays and Fridays), which is a standard diet in larval lamprey studies. All experimental groups were maintained under identical feeding conditions. Future studies should consider that larval sea lampreys, which are filter feeders, obtained and kept in different parts of the world might be exposed to different microbial communities present in the water.

Control sham injured animals were processed as above by performing the dorsal incision of the body wall but without performing the SCI surgery. Control non injured animals and sham controls were always processed in parallel to animals with a SCI.

### 2.2 Gentamicin treatment

Larvae with a complete SCI (n=16) were treated with the broad-spectrum antibiotic gentamicin. Gentamicin (Sigma, ref. G1264) was dissolved in the 200 mL of freshwater where the animals were left after surgery at a concentration of 15 µg/ml. Lampreys were always injured on a Monday and the 200 mL of water with new gentamicin were replaced each Monday, Wednesday and Friday of every week (3 treatments per week) during the first 5 weeks post-injury. After this 5-week gentamicin treatment, 6 larvae were processed for the gut microbiome analysis (see section 3.3) and 10 larvae were kept in freshwater until they were 10 weeks post-injury for the analysis of axon regeneration (see section 3.4). Control non-treated lampreys with a complete transection SCI (n = 10) were processed in parallel to gentamicin-treated lampreys but were only maintained in 200 ml of freshwater that was also replaced 3 times per week as with treated lampreys. Control and treated lampreys were fed with baker’s yeast as indicated above.

### 2.3 Analysis of gut microbial communities by 16S rRNA Gene Sequencing

At 5 or 8 weeks post-surgery, six lamprey larvae per group (n = 42 in total) were euthanized and dissected to collect whole individual intestinal content samples, which were immediately stored at −20 °C until further processing (gentamicin treated larvae were only analysed at 5 weeks post-surgery). Total DNA was extracted using the E.Z.N.A Tissue DNA kit (Omega Biotek) following the manufacturer’s protocol. The V3–V4 hypervariable region of the 16S rRNA gene was amplified with primers Bakt_341F (5′-CCTACGGGNGGCWGCAG-3′) and Bakt_805R (5′-GACTACHVGGGTATCTAATCC-3′), as described by Herlemann et al. (2011). PCR products were verified by agarose gel electrophoresis, purified with Mag-Bind RXNPure Plus magnetic beads (Omega Bio-tek), and indexed using Nextera XT adapters (Illumina). Sequencing was performed on an Illumina MiSeq platform with 2×300 bp paired-end reads. Amplicons were randomly grouped into 3 replicates (R1, R2 and R3), each containing 2 larval samples of the same experimental group, to assess reproducibility.

Raw reads were quality filtered and trimmed using PRINSEQ, removing sequences with ambiguous bases, low complexity, or average Phred scores below 25. Taxonomic classification was performed with Kraken2 (Wood et al., 2019) against the SILVA 16S rRNA reference database, and relative abundances were computed at different taxonomic levels in QIIME2 (Boylen et al., 2019). Alpha- and beta-diversity indices were calculated, and Bray-Curtis dissimilarities were visualized by Principal Coordinates Analysis (PCoA). Statistical significance of microbiota community differences among groups was assessed using PERMANOVA (adonis function, 9,999 permutations) implemented in the vegan v2.6-4 package in R.

### 2.4 Anti-CCK-8 immunofluorescence, confocal imaging and quantification of axon regeneration

The spinal cord region between the 4^th^ and 6^th^ gill openings of 10 weeks post-lesion (wpl) animals (with or without gentamicin treatment) was dissected out and fixed by immersion in 4% paraformaldehyde in 0.05 M Tris-buffered saline (TBS) pH 7.4 for 6 h at 4 ºC. The spinal cord was then rinsed in TBS, cryoprotected in 30% sucrose in TBS overnight at 4ºC, embedded in Neg-50 (Epredia), frozen in liquid nitrogen, and cut on a cryostat (18 µm transverse sections). Sections were mounted on Superfrost® Plus glass slides (Epredia). Then, they were incubated with a purified rabbit polyclonal anti-sea lamprey cholecystokinin (CCK)-8 antibody (dilution 1:200; see Sobrido-Cameán et al., 2020c) at RT overnight. After washes in TBS, the sections were incubated for 1h at RT with a Cy3-conjugated goat anti-rabbit antibody (1:500; Millipore; Cat# AP132C; RRID: AB_92489). Antibodies were always prepared in a solution containing 15% normal goat serum and 0.2% Triton. Sections were then rinsed in TBS and distilled water and mounted with Mowiol (Sigma). The specificity of the purified anti-CCK-8 antibody was previously confirmed by ELISA and by comparison with *in situ* hybridization experiments for the detection of the CCK mRNA (Sobrido-Cameán et al., 2020c).

A Stellaris 8 confocal microscope (Leica) was used to acquire images of the spinal cord sections using a 20x objective. Optical sections were taken at steps of 0.7 μm along the z-axis. Collapsed projections of spinal cord sections were obtained with the LAS X software (Leica). Figures were generated using Adobe Photoshop 2024 (Adobe). Confocal photomicrographs were converted to B&W in the figures to facilitate viewing.

CCK-immunoreactive (ir) regenerated axon profiles were quantified in confocal photomicrographs of the 18 µm transverse sections using the Feature J plugin of the Fiji (Schindelin et al., 2012) software (see Fernández-López et al., 2014). The images from control and 10 wpl animals were taken with the same confocal microscope settings and using the same software parameters. A threshold was determined to obtain the most accurate images when converting them to binary B&W images for axon profile quantification. We quantified the number of CCK-ir axon profiles in 1 out of every 4 consecutive spinal cord sections starting at the level of the 6th gill opening and moving rostrally. 9 sections were analyzed per animal covering 648 µm of spinal cord below the site of injury. The mean number of CCK-ir axon profiles per section was obtained for each animal based on the quantification of the 9 sections (each data point in the graphs represents one larva).

Statistical analyses were carried out with Prism 9 (GraphPad). We first checked the data (mean number of regenerated CCK-ir axon profiles per section) for normality using the D′Agostino-Pearson test. To assess any significant differences (with p ≤0.05) between the control non-treated and the gentamicin-treated groups, we used a two-tailed unpaired Student’s t-test, assuming the data followed a normal distribution based on the results from the D′Agostino-Pearson test. Each experimental group, consisting of 10 animals, was obtained from 2 separate batches of animals. Within each batch, control and 10 wpl animals were processed in parallel. 1 sample from the control group was lost during the spinal cord dissection at the end of the 10-week period.

## 3 Results

### 3.1 A complete spinal cord injury induces changes in the gut microbiome community of mature larval sea lampreys

Analysis of taxonomic composition revealed marked differences in the sea lamprey gut microbial composition among treatments and recovery times (Figure 1). Control and sham control groups displayed communities enriched in *Legionellaceae, Bradyrhizobiaceae* and *Planctomycetaceae*, whereas animals with a complete SCI were dominated by *Legionellaceae*. In contrast, antibiotic-treated injured animals showed a profoundly altered profile, with near-complete replacement of microbiota diversity by *Bradyrhizobiaceae* (see below).

**Table 1.** Median (± SD) relative abundance (%) of bacterial families across treatments and recovery times. Median and SD values were calculated from three biological replicates per group (C, control; SC, sham control; SCI, spinal cord injury; SCI+Ab, spinal cord injury with antibiotic treatment).

| Family Name | C 5W | C 8W | SC 5W | SC 8W | SCI 5W | SCI 8W | SCI+Ab 5W |
| --- | --- | --- | --- | --- | --- | --- | --- |
| *Legionellaceae* | $0.328 \pm 0.036$ | $0.530 \pm 0.010$ | $0.088 \pm 0.006$ | $0.598 \pm 0.049$ | $0.815 \pm 0.016$ | $0.590 \pm 0.006$ | $0.002 \pm 0.000$ |
| *Bradyrhizobiaceae* | $0.311 \pm 0.029$ | $0.130 \pm 0.009$ | $0.733 \pm 0.000$ | $0.080 \pm 0.002$ | $0.048 \pm 0.000$ | $0.181 \pm 0.009$ | $0.986 \pm 0.000$ |
| *Planctomycetaceae* | $0.221 \pm 0.002$ | $0.156 \pm 0.008$ | $0.039 \pm 0.004$ | $0.103 \pm 0.052$ | $0.007 \pm 0.001$ | $0.021 \pm 0.002$ | $0.000 \pm 0.000$ |
| *Hyphomicrobiaceae* | $0.027 \pm 0.001$ | $0.095 \pm 0.003$ | $0.022 \pm 0.001$ | $0.049 \pm 0.006$ | $0.024 \pm 0.010$ | $0.051 \pm 0.000$ | $0.001 \pm 0.000$ |
| *Microbacteriaceae* | $0.025 \pm 0.004$ | $0.017 \pm 0.001$ | $0.014 \pm 0.000$ | $0.020 \pm 0.001$ | $0.016 \pm 0.003$ | $0.029 \pm 0.002$ | $0.000 \pm 0.000$ |
| *Gemmataceae* | $0.003 \pm 0.001$ | $0.006 \pm 0.001$ | $0.004 \pm 0.000$ | $0.024 \pm 0.003$ | $0.019 \pm 0.006$ | $0.034 \pm 0.001$ | $0.000 \pm 0.000$ |
| *Methylobacteriaceae* | $0.005 \pm 0.001$ | $0.002 \pm 0.000$ | $0.024 \pm 0.002$ | $0.005 \pm 0.000$ | $0.020 \pm 0.008$ | $0.017 \pm 0.000$ | $0.006 \pm 0.000$ |
| *Xanthobacteraceae* | $0.000 \pm 0.000$ | $0.026 \pm 0.001$ | $0.000 \pm 0.000$ | $0.051 \pm 0.010$ | $0.000 \pm 0.000$ | $0.000 \pm 0.000$ | $0.000 \pm 0.000$ |
| *No family* | $0.024 \pm 0.003$ | $0.010 \pm 0.001$ | $0.010 \pm 0.000$ | $0.006 \pm 0.003$ | $0.001 \pm 0.000$ | $0.006 \pm 0.000$ | $0.000 \pm 0.000$ |
| *Brucellaceae* | $0.003 \pm 0.001$ | $0.003 \pm 0.000$ | $0.016 \pm 0.002$ | $0.019 \pm 0.002$ | $0.009 \pm 0.000$ | $0.002 \pm 0.000$ | $0.000 \pm 0.000$ |
| *Other* | $0.053 \pm 0.003$ | $0.026 \pm 0.004$ | $0.049 \pm 0.000$ | $0.045 \pm 0.004$ | $0.040 \pm 0.003$ | $0.068 \pm 0.001$ | $0.004 \pm 0.001$ |

**Figure 1.**
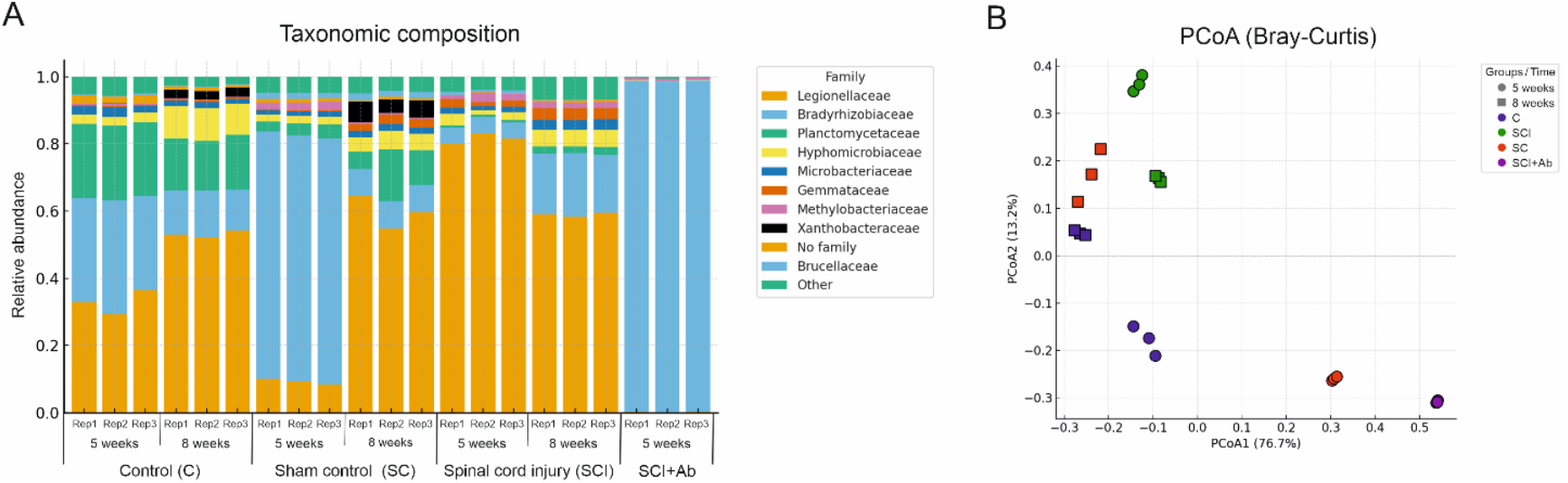
Taxonomic composition and beta-diversity of intestinal microbiota across treatments and recovery times. (**A**) Relative abundance of bacterial families in each experimental group: control (C), sham control (SC), spinal cord injury (SCI), spinal cord injury treated with antibiotic (SCI+Ab), time points (5 and 8 weeks); and biological replicates (Rep1, Rep2 and Rep3). (**B**) Principal coordinate analysis (PCoA) based on Bray-Curtis differences. The first principal coordinate (PCoA1) explains 76.7% of the variance, while the second (PCoA2) accounts for 13.2%. The plot illustrates significant differences among groups (PERMANOVA, pseudo-F = 36.89, p = 0.001) and time (pseudo-F = 28.14, p = 0.017), with C, SC, and SCI groups tending toward convergence at 8 weeks.

PCoA based on Bray-Curtis dissimilarities confirmed significant effects of the treatments (PERMANOVA, pseudo-F = 36.89, p = 0.001) and a global effect of time (pseudo-F = 28.14, p = 0.017), with control, sham control and injured groups tending toward convergence at 8 weeks. This temporal effect was further reflected in average Bray-Curtis distances, which decreased from 0.42 ± 0.05 at 5 weeks to 0.29 ± 0.04 at 8 weeks across these groups.

Control and sham control groups did not differ significantly at either 5 weeks (pseudo-F = 12.37, p = 0.087) or 8 weeks (pseudo-F = 9.00, p = 0.631). However, sham control animals exhibited an initial shift in intestinal microbiota composition (reflected by an expansion of *Bradyrhizobiaceae*), with an average Bray-Curtis distance of 0.46 ± 0.02 from controls at 5 weeks, which decreased to 0.15 ± 0.05 by 8 weeks.

In contrast, injured animals differed significantly from controls (pseudo-F = 22.18, p = 0.004), a divergence associated with reduced *Planctomycetaceae* and *Xanthobacteraceae* and concomitant expansion of *Legionellaceae*. These differences were particularly evident at 5 weeks (p = 0.012), whereas by 8 weeks the difference was no longer significant (p = 0.281). Pairwise comparisons between injured animals and sham controls also revealed significant differences at 5 weeks (p = 0.024), but not at 8 weeks (p = 0.317). Together, these findings indicate that after an initial shift associated with the SCI (expansion of *Legionellaceae*), a temporal effect on microbiota structure was observed, indicating convergence and partial recovery toward a control-like community.

### 3.2 A gentamicin treatment causes changes in the gut microbiome community of injured lampreys

Gentamicin is an aminoglycoside broad spectrum antibiotic that works by binding the 30S subunit of the bacterial ribosome, negatively impacting protein synthesis. Metabarcoding analysis of SCI larvae treated with gentamicin (SCI+Ab) confirmed that antibiotic exposure for 5 weeks caused a severe disruption of the gut microbiota, as community composition in these animals was restricted almost exclusively to *Bradyrhizobiaceae* members (Figure 1). Thus, SCI+Ab samples were highly distant from all other groups (mean Bray-Curtis distance ≥ 0.85) and showed significant divergence in all pairwise comparisons (p ≤ 0.02), confirming that gentamicin treatment induced a profound disruption of the microbiota. This result correlates with the failure to recover bacterial isolates from the fecal samples of this group by culture-based microbiological methods (data not shown).

Specifically, the SCI+Ab group differed from SCI animals, with an average Bray-Curtis distance of 0.95 ± 0.01 (p = 0.02), underscoring the strong divergence between antibiotic-disrupted and lesion-associated microbiotas.

### 3.3 Gentamicin-induced microbiome dysbiosis does not affect the spontaneous axon regeneration of descending cholecystokinergic neurons

We have recently shown that descending cholecystokinergic axons of the sea lamprey regenerate spontaneously (81% of the number of CCK-ir axon profiles of control non-injured animals) at the level of the 6^th^ gill 10 weeks after a complete transection SCI at the level of the 5^th^ gill (González-Llera et al., 2024). This descending neuropeptidergic system offers a model of interest to identify new signalling systems regulating spontaneous axon regeneration because it provides and axon system in which axon regeneration can be promoted or inhibited via drug or genetic manipulations. Since all spinal cord cholecystokinergic axons of the larval sea lamprey spinal cord come from brainstem neurons of the caudal rhombencephalon and not from intraspinal neurons (Sobrido-Cameán et al., 2020c; González-Llera et al., 2024), we can evaluate the impact of a specific treatment on axon regeneration by means of a simple immunofluorescence protocol and by quantifying the number of CCK-ir regenerated axon profiles in transverse sections of the cord, without the need to use neuronal tracers.

Thus, we used an antibody generated against the sea lamprey CCK-8 peptide (Sobrido-Cameán et al., 2020c) to study the effect of a gentamicin treatment on the regeneration of descending cholecystokinergic axons. We treated the animals with gentamicin during the first 5 wpl because this is a crucial period in the process of spontaneous axon regeneration in lampreys [axon retraction predominates in the first wpl, axon regrowth starts to predominate after the second wpl and some axons begin to cross the injury site by 4-5 wpl; (Zhang et al., 2005)]. Surprisingly, and despite the drastic changes observed in the gut microbiome (see section 3.2), the 5-week gentamicin treatment did not cause a change in the number of regenerated CCK-ir axon profiles present at the level of the 6^th^ gill 10 weeks after a complete transection SCI at the level of the 5^th^ gill (Figure 2).

**Figure 2.**
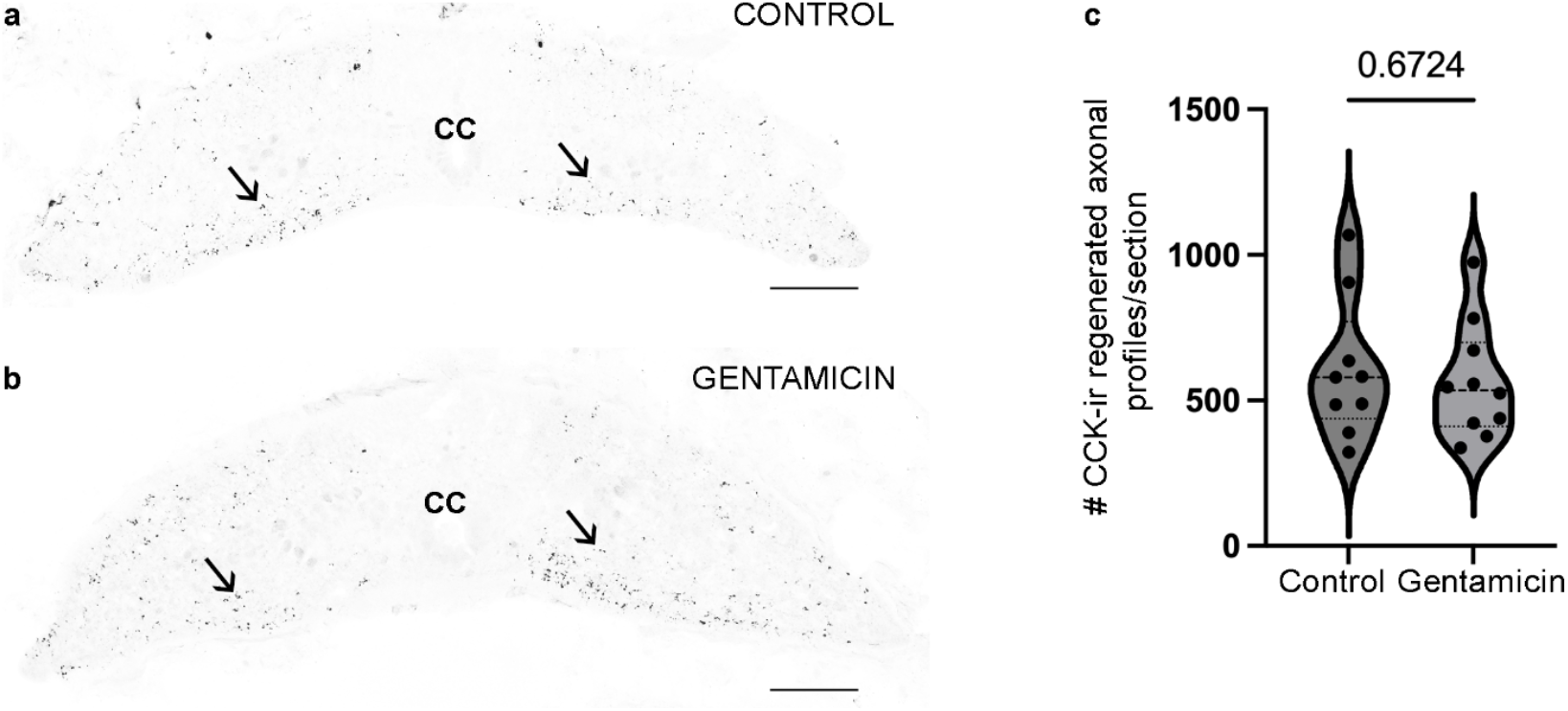
A 5-week gentamicin treatment does not affect the spontaneous regeneration of descending cholecystokinergic axons. Transverse spinal cord sections of control non-treated (a) and gentamicin-treated (b) 10 wpl larvae (level of the 6th gill) showing similar numbers of CCK-ir regenerated axon profiles (arrows) [see graph in c, each dot represents 1 animal; control non-treated (n= 9): 606.1 ± 80.05 CCK-ir regenerated axon profiles; gentamicin-treated (n = 10): 562.9 ± 62.50 CCK-ir regenerated axon profiles]. Cc: central canal. Scale bars: 75 µm.

## 4 Discussion

Our results revealed that a complete spinal cord transection in larval sea lampreys causes an initial shift in the composition of microbial communities of the gut, which is reflected as reduced presence of *Planctomycetaceae* and *Xanthobacteraceae* and concomitant expansion of *Legionellaceae* as compared to controls. Sham controls also showed an initial shift in the composition of microbial communities associated with a (non-significant) expansion of *Bradyrhizobiaceae* members as compared to non-injured controls. Thus, spinal cord damage specifically alters intestinal microbial composition (expansion of *Legionellaceae)*, which is independent of the damage to the body wall tissues that was also performed in sham controls. Our results, together with previous data in rodent models and human patients (Kigerl et al., 2016; Jogia and Ruitenberg, 2020; Du et al., 2021; Jing et al., 2021, 2023; Kong et al., 2023; Wilson et al., 2023), indicate that gut microbial disruption after a SCI is a shared characteristic of regenerating (i.e. lampreys) and non-regenerating (i.e. mammals) vertebrates.

Based on our results on the changes in microbial communities in lampreys, we hypothesized that the initial expansion in *Legionellaceae* (at 5 weeks post-injury) could be positively correlated with the process of spontaneous and successful axon regeneration that occurs after a complete SCI in lampreys (e.g., giant descending neurons: Jacobs et al., 1997; descending serotonergic neurons: Cornide-Petronio et al., 2011; descending neuropeptidergic neurons: González-Llera et al., 2022, 2024). Unexpectedly, our treatments of injured lampreys with the broad-spectrum antibiotic gentamicin, which caused a complete disruption of the gut microbial communities (i.e. loss of *Legionellaceae* and expansion of *Bradyrhizobiaceae*), indicate that the intestinal expansion of *Legionellaceae* is dispensable for spontaneous regeneration of descending cholecystokinergic axons. The degree of regeneration of descending cholecystokinergic axons in larval sea lampreys correlates with functional swimming recovery, as regenerated axons re-establish synaptic contacts caudal to the lesion site (González-Llera et al., 2024). Thus, the absence of differences in CCK-ir axon regeneration between control and antibiotic-treated injured animals suggests that behavioural recovery would be preserved despite profound microbiome disruption, although this should be experimentally confirmed in the future.

On the other hand, the gentamicin treatment in injured lampreys caused a loss of most families and an expansion of *Bradyrhizobiaceae*, a family that is also present in lampreys after a complete transection SCI without an antibiotic treatment. So, future studies could attempt to perturb the intestinal presence of members of this family specifically to determine their possible influence on the spontaneous regeneration of cholecystokinergic axons observed in both injured controls and gentamicin-treated injured animals. Moreover, it would be also of interest to determine whether the regeneration of other descending systems (e.g. giant descending neurons or descending serotonergic neurons) is specifically influenced by the changes in gut microbial communities observed in lampreys with a complete SCI (i.e. expansion of *Legionellaceae*).

Overall, these findings demonstrate that while SCI induces transient gut microbiota shifts in larval sea lampreys, such changes do not modulate spontaneous axon regeneration. This suggests that the regenerative capacity of lamprey neurons is robust to microbial perturbations, distinguishing them from mammalian systems where gut microbiota alterations and subsequent microbiota-immune-neuronal interactions may influence recovery. Future studies dissecting specific microbial–immune-neuronal interactions may help clarify how host-microbe relationships evolved alongside regenerative abilities in vertebrates.

## 6 Author Contributions

Conceptualization, G.N.S.-D., M.B., A. B.-I.; investigation, L.G.-L., G.N.S.-D., A. V., N. B., M.B.; writing—original draft preparation, L.G.-L., M. B., A.B.-I.; writing—review and editing, G.N.S.-D., A. V., N. B.; funding acquisition, G.N.S.-D., M.B., A. B.-I..

## 8 Acknowledgments

We would like to thank the *Servizo de Microscopía* of the Universidade de Santiago de Compostela and Dr. Mercedes Rivas Cascallar for confocal microscope facilities and technical help.

## 10 Data Availability Statement

All sequencing reads have been deposited in the NCBI Sequence Read Archive (SRA) under BioProject accession number PRJNA1387090.

Datasets generated for this study can be found in the following repositories:

DNA sequences: Genbank, accession numbers F234391-F234402.

Phylogenetic data, including alignments: TreeBASE, accession number S9123.

